# Assessing Behavioral and Neural Correlates of Change Detection in Spatialized Acoustic Scenes

**DOI:** 10.1101/2024.12.03.626637

**Authors:** Katarina C. Poole, Drew Cappotto, Vincent Martin, Jakub Sztandera, Maria Chait, Lorenzo Picinali, Martha Shiell

**Author notes:** Corresponding author: Katarina C. Poole.

## Abstract

The ability to detect changes in complex auditory scenes is crucial for human survival, yet the neural mechanisms underlying this process remain elusive. This study investigates how the presence and location of sound sources impacts active auditory change detection as well as neural correlates of passive change detection. We employed stimuli designed to minimize semantic associations while preserving naturalistic temporal envelopes and broadband spectra, presented in a spatial loudspeaker array. Behavioral change detection experiments tasked participants with detecting new sources added to spatialized and non-spatialized multi-source auditory scenes. In a passive listening experiment, participants were given a visual decoy task while neural data were collected via electroencephalography (EEG) during exposure to unattended spatialized scenes and added sources.

Our behavioral experiments (N = 21 and 21) demonstrated that spatializing sounds facilitated change detection compared to non-spatialized presentation, but that performance declined with increasing number of sound sources and higher hearing thresholds at high frequencies, exclusively in spatialized conditions. Slower reaction times were also observed when changes occurred from above or behind the listener, exacerbated by a higher number of sources. EEG experiments (N = 32 and 30), using the same stimuli, showed robust change-evoked responses. However, no significant differences were detected in our analysis as a function of spatial location of the appearing source.

## 1. INTRODUCTION

The ability to rapidly detect changes in complex acoustic environments is crucial for survival and for effective interactions with the outside world. Human listeners demonstrate a remarkable capacity to parse intricate auditory scenes into distinct auditory objects and streams (Bregman, 1994; Shinn-Cunningham, 2008; Snyder and Alain, 2007) and in detecting any changes in the auditory scene in the form of appearing and disappearing sound sources (Aman et al., 2021; Constantino et al., 2012; de Kerangal et al., 2021; Demany et al., 2017; Gregg and Samuel, 2008; Irsik et al., 2016; Pavani and Turatto, 2008; Sohoglu and Chait, 2016a). This rapid detection of changes in ongoing acoustic input serves the listener as a 360° ‘early warning’ system, supporting situational awareness and allowing the listener to quickly adjust their behavior without relying on visual cues. Those who struggle with detecting changes in their auditory environment — such as individuals with hearing impairments or age-related decline in auditory processing (de Kerangal et al., 2021)— may face challenges in staying alert to their surroundings, impacting their quality of life and safety (Ju et al., 2022). Therefore, it is important to understand the auditory mechanisms underlying auditory change detection and how we can effectively assess this ability in a listener.

Various factors can influence a listener’s ability to detect a scene change. One notable finding is that listeners tend to be more accurate and quicker at detecting sounds that appear (onsets) than sounds that disappear (offsets) from a given auditory scene (Constantino et al., 2012; Huron, 1989; Pavani and Turatto, 2008; Sohoglu and Chait, 2016a). This perceptual asymmetry may stem from differences in neural responses triggered by auditory onsets versus offsets, where auditory onsets tend to elicit more pronounced neural responses (Pantev et al., 1996; Phillips et al., 2002; Pratt et al., 2008). In addition, the role of attention in change detection remains an ongoing subject of inquiry, with studies suggesting that selective attention can influence the perception of auditory objects and streams (Shinn-Cunningham, 2008), where attention-dependent listener tasks decrease the prevalence of change “deafness”. Eramudugolla et al., (2005) showed that when a listener had to identify the disappearance of a sound in a scene of multiple concurrent naturalistic sounds, directing attention to the potential change remarkably increased detection accuracy. Other factors such as the predictability of an auditory scene (Aman et al., 2021; de Kerangal et al., 2021; Sohoglu and Chait, 2016b) and listener expertise (Vanden Bosch der Nederlanden et al., 2020) have all been shown to modulate auditory change detection.

An important factor to consider is that real-world acoustic environments vary in density as a function of the number of discrete sound objects present at a given scene. Research using common environmental sounds has shown that when concurrent background streams are more distinct and segregable, with a larger acoustic spread, detection is facilitated (Gregg and Snyder, 2012). Other studies, using concurrent pure tone-pip sequences, demonstrated that as the number of background streams increases, detecting both appearing and disappearing sounds becomes more challenging (Aman et al., 2021; Sohoglu and Chait, 2016a). One key cue for separating auditory streams is the spatial location of each individual sound source.

Spatial cues are known to play a crucial role in perceptual grouping, allowing listeners to separate and track individual sound sources within the acoustic environment, even when they are not behaviorally relevant (Snyder and Alain, 2007). Spatial separation of sources should lead to improved change detection and has indeed been demonstrated by recent studies showing that it can significantly affect task performance in alternate-speaker and spatial speech-in-noise tasks (Salorio-Corbetto et al., 2022; Uhrig et al., 2022). Yet the role of spatialization in auditory change detection has yielded mixed results. Aman et al. (2021) found that spatialization enhanced change detection of an appearing sound but not a disappearing sound. In contrast, Eramudugolla et al. (2005) observed detection of a disappearing sound source was enhanced when the sounds were presented at distinct spatial locations than when co-located, but only when the listener’s attention was nondirected. Furthermore, Gregg and Samuel (2008) demonstrated an impairment to change detection with spatialization and proposed that this could be due to an attentional spatial spotlight focusing the listener away from the change (Best et al., 2006). Subsequent research has highlighted the role of the attentional spotlight in the context of gaze-directed attention. Pomper and Chait (2017) demonstrated faster reaction times for detecting deviant tones when participants fixated their gaze on the target location rather than a distractor location. Similarly, Best et al. (2023) showed that gaze fixation on the location of a target talker improves speech intelligibility in multi-talker environments. Overall, these findings suggest that spatialization exerts a nuanced effect on a listeners’ ability to detect scene changes. On one hand, it enhances detection by perceptually segregating sound sources, improving focus on the target. On the other, it can impair detection—for changes occurring outside the gaze angle—by diverting attention away from the change.

To elucidate the cognitive processes underlying auditory change detection, neuroimaging studies using electroencephalography (EEG) and magnetoencephalography (MEG) have identified change-evoked responses in the event-related potential (ERP) that suggest auditory change detection is an automatic and multi-staged process (Puschmann et al., 2013; Snyder et al., 2015; Sohoglu and Chait, 2016a). Early ERP components show an increase in amplitude in the auditory cortex for both detected and undetected scene changes when compared to no-change conditions (e.g. the Nb component 30-40 ms after change onset). Later-stage neural responses (e.g. >100 ms) associated with higher-level cognitive processes have instead been shown to modulate as a function of successfully detected changes (Gregg et al., 2014; Gregg and Snyder, 2012; Puschmann et al., 2013; Snyder et al., 2015; Sohoglu and Chait, 2016a). For example, Sohoglu & Chait (2016) used magnetoencephalography (MEG) to examine the neural dynamics of change responses in complex auditory scenes, observing increased and sustained activity in the auditory and parietal cortices emerging around 100-300 ms after the appearance of a new scene source, with amplitude increases as a function of detected vs. undetected scene changes. This suggests that, while scene changes are detected automatically early in the primary auditory cortex, separable mechanisms are responsible for upstream feature processing related to their perception.

Much of the existing behavioral and neuroimaging literature has employed non-spatialized and relatively simple stimuli such as tone pips or noise bursts. These do not fully capture the spectrotemporal complexity of real-world auditory scenes or contain stimuli known to modulate responses based on semantic meaning or familiarity. The spatialization has also been limited to simple left-right spatialization protocols, leaving the behavioral and neural correlates of change detection in spatialized scenes a largely underexplored topic. Here we investigate the behavioral effects and neural correlates underlying change detection in spatialized acoustic scenes in four experiments. "Chimeric" sounds (Smith et al., 2002) were designed to preserve the spectrotemporal complexity of real-world auditory environments, and retain broadband frequency content for binaural and monaural spatial cues, all while minimizing semantic associations that have been shown to modulate change response features in previous studies (Gregg et al., 2014; Ozmeral and Menon, 2023).

We investigated whether behavioral change detection is affected by the spatialization of sound sources by assessing reaction times and detection accuracy to an appearing sound source; hypothesizing that spatial separation of sources would facilitate change detection (Experiment 1). We assessed automatic change detection with a passive listening paradigm whilst measuring neural responses using electroencephalography (EEG, Experiments 3 and 4). This was to identify whether we would observe neural responses consistent with previous literature on auditory change detection despite a more complex and spatialized stimulus. Lastly, we examined how behavioral and neural responses are affected by the location of the appearing sound source (Experiments 2, 3 and 4). We hypothesized that a bias towards the front as reported in previous work (Best et al., 2023; Pomper and Chait, 2017) may diminish sensitivity to sources above and/or behind the listener. This reduced sensitivity is expected to result in poorer detection performance, as well as delayed and weaker neural change responses. By probing change detection at different locations in spatialized acoustic scenes with acoustically rich yet unfamiliar sounds, our study provides novel insights into the processes supporting auditory scene analysis and change detection in naturalistic spatialized auditory scenes.

## 2. MATERIAL AND METHODS

### 2.1 Stimuli

We aimed for stimuli that had minimal semantic associations but still preserved properties from real-world sounds, specifically complex temporal envelopes and spectra. We employed a method for creating so-called “chimeric” sounds (Smith et al., 2002), by extracting the temporal envelope (low-pass filtered at 10 Hz) from modulator sounds (e.g., animal vocalizations, babble speech, environmental recordings) and applying it to synthesized carrier sounds. The carrier sounds were characterized by diverse fundamental frequencies, noise, and harmonic content (see Figure 2.1A-B), and all had a broad spectrum to maintain localization cues (Blauert, 1997). Chimeras underwent 300 Hz Butterworth high-pass filtering and loudness normalization (-24 LUFS). They were trimmed to nine seconds in length and manually edited to reduce amplitude fluctuations. Due to the variations between chimeras, we performed pilot testing in which all chimeras could be an appearing sound source and background source during the experiment. From the behavioral data we then selected five chimeras (associated with best performance) to then designate as “appearing sources” in further experiments, whilst the other ten served as "background sources". See the code and data repository to generate the spatialized stimuli at: *github.com/Audio-Experience-Design/ANTHEA_spatialized_change_detection*.

**Figure 2.1.**
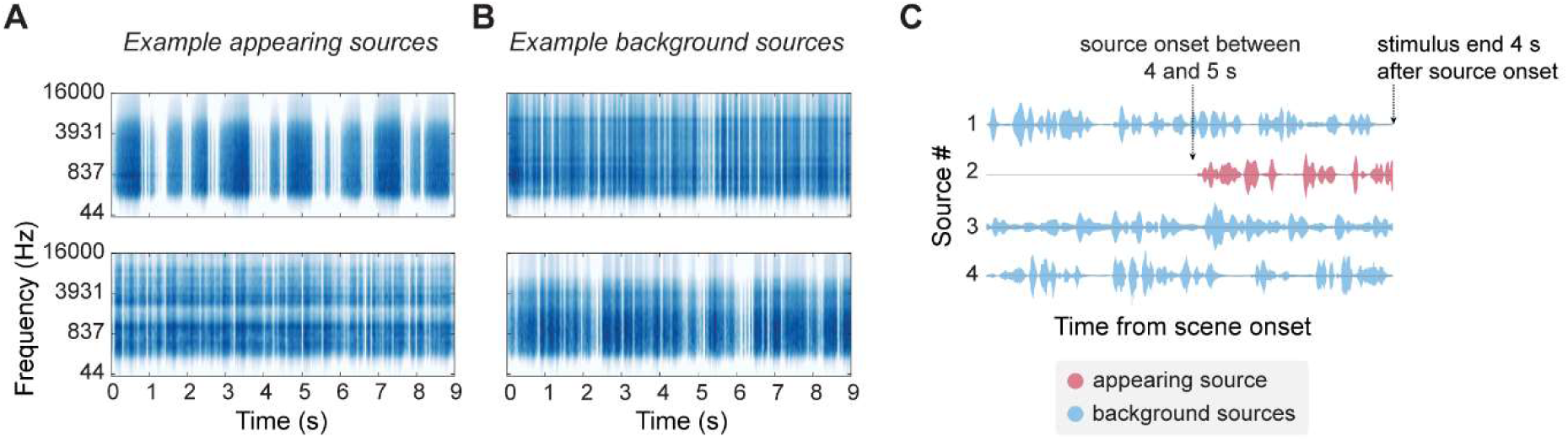
Acoustic stimuli and scene structure. ERB-weighted spectra of two examples of appearing sources (A) and two examples of background sources (B). (C) Example schematic of the acoustic scenes used in Experiments 1 and 2 with concurrent sources and an appearing sound source 4 to 5 seconds from the onset of the acoustic scene. All sources overlapped in frequency space and the whole stimulus ended 4 seconds after appearing source onset. In Experiment 3 and 4 the appearing source onset was fixed at 4 seconds from the scene start and persisted for 3 seconds. In catch trials there was no appearing sound source and the whole acoustic scene lasted between 8 and 9 seconds for Experiment 1 and 2, and 7 seconds in Experiments 3 and 4.

### 2.2 Participants

Experiment 1 included 21 paid participants; (12 males, mean age 30 years, standard deviation 13 years). Experiment 2 included 21 paid participants (13 males, mean age 29, standard deviation 12 years). Experiment 3 included 32 paid participants (22 males, mean age 30 years, standard deviation 13 years). Experiment 4 included 30 paid participants (21 males, mean age 29 years, standard deviation 9 years) with eight excluded in Experiment 3 due to technical issues. When participants took part in more than one of the four experiments, data collection was undertaken on different days and experiments 3 or 4 (EEG) were performed first, to avoid any behavioral bias that could influence the neural recordings. All participants reported normal hearing and no neurological disorders. Experimental procedures for Experiments 1 to 3 were approved by the Imperial College London ethics committee (SETREC number: 6533981) and Experiment 4 was approved by the UCL Research Ethics Committee (Project ID Number: 1490/011). Written informed consent was obtained from each participant.

### 2.3 Procedure

Stimuli were presented via a 31-loudspeaker array (see Figure 2.2A-B) in a semi-anechoic room for Experiments 1,2 and 3. In Experiment 4, an 8-loudspeaker setup was used (see Figure 2.2C-D and section 2.3.4 for details on the phantom source rendering). For all experiments, the average sound pressure level at the participant position was 22 dBA when all loudspeakers were silent. During the stimulus presentation, the loudspeaker arrays were calibrated so that each sound scene had an average level between 68 dBA and 70 dBA at the location of the listener, depending on the number of sound sources in the scene (4 to 8). The participant’s head was always positioned at the center of the loudspeaker array using an adjustable chair. Max/MSP software (Cycling74) managed stimulus delivery via an Antelope Audio Orion 32 sound card, triggered by custom Python scripts (Python Software Foundation) that also provided text prompts and visual stimuli to a screen placed inside the apparatus and in front of the participant. In all experiments, participants were briefly trained prior to beginning the task. Experiment 1 was 40 minutes in length and Experiments 2, 3 and 4 were 60 minutes in length, with participants given a break every 10 minutes.

**Figure 2.2.**
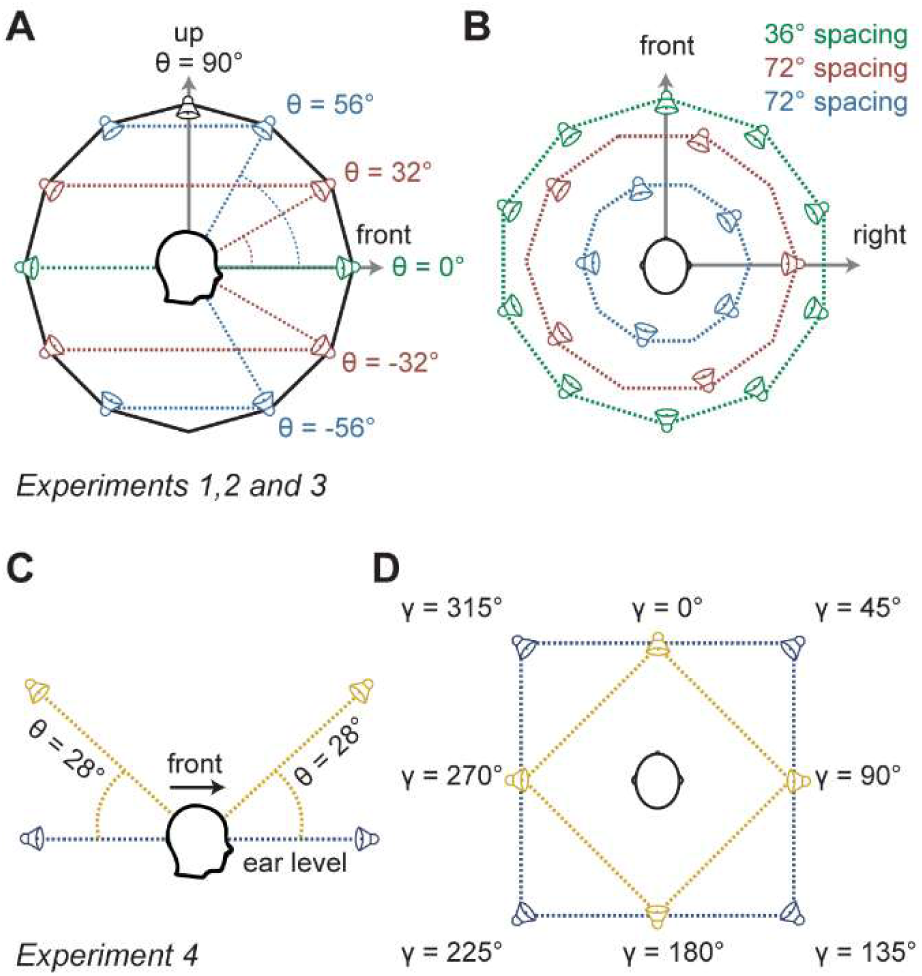
Loudspeaker array schematics. (A,B) Spherical loudspeaker array layout used for Experiments 1, 2 and 3. The participant was situated at the center of the sphere. The array consisted of 31 loudspeakers organized across 5 planes at varying elevations and separations (A). Additionally, a speaker (depicted in black) was placed atop the sphere directly above the participant. (C,D) Loudspeaker array layout used for Experiment 4. The array consisted of 8 loudspeakers organized across 2 distinct planes at varying elevations and 90° separations (C). The two planes are offset in azimuth of 45°, so all 8 speakers are equally spaced of 45° in azimuth with the participant situated in the center.

#### 2.3.1 Experiment 1 (Behavioral)

In Experiment 1 we tested the effects of spatialization and the number of concurrent sound sources on the behavioral performance in the detection of an appearing sound source. We measured the participants’ reaction time and accuracy and varied the number of sources in the scene (5 conditions: between 4-8 sources, including the appearing source). To test the effect of spatialization, scenes were presented either with each source assigned to a random, unique location (spatialized scene condition), or every loudspeaker playing all sources simultaneously (non-spatialized scene condition). Trials were presented in four blocks. Spatialization conditions (spatialized or non-spatialized scenes) were tested in separate blocks, with the first two blocks belonging to one of these conditions, and the order of these conditions randomized between participants.

The number of sources in the scene varied across trials within each block in a random order. Within a block, an inter-trial-interval of 1 to 2 s was used. Half of the trials contained a change in the form of an appearing source, the onset of which occurred between 4 and 5 s from scene onset and ended after an additional 4 s (see Figure 2.1C). Appearing sources were presented pseudo-randomly from any of the 31 speakers, with 10 repeats for each number of sources in the scene (five conditions), for a total of 200 trials. The locations of the background sources were also restricted, such that they appeared in pseudo-randomly selected locations to ensure a balanced distribution around the listener. In trials with no appearing sound source, the no change condition, the scenes lasted between 8 and 9 s. Participants were instructed to press a button as quickly as possible when they heard an appearing sound source within a trial, and that not all trials would contain appearing sources.

Since spatialization benefits may be influenced by a participant’s high-frequency hearing, we assessed participants’ pure-tone thresholds for frequencies 2, 4, 8, 12 and 16 kHz. We employed a 2-down 1-up adaptive staircase method. The staircase started at 20 dBA, changed by 8 dB for the first three reversals and then 2 dB for the next three reversals (limited at 85 dBA), stopping after six reversals. The mean over the last three reversals was taken as the threshold for each frequency at each ear individually. Pure tones were 1s in length with a 90 ms cosine on/off ramp, and were presented at a random inter-stimulus interval between 1 and 3 s. The pure-tone assessment occurred at the beginning of the experiment and lasted approximately 20 minutes (10 minutes for each ear). The participants were sat within the loudspeaker array and the stimuli were delivered through Sennheiser HD 599E headphones.

#### 2.3.2 Experiment 2 (Behavioral)

In Experiment 2 we examined how the location of an appearing sound source may influence detection performance. We used the same paradigm as in Experiment 1 but restricted the locations of the appearing source to five location conditions: front (azimuth = 0, elevation = 0), above (azimuth = 0, elevation = 90), back (azimuth = 0, elevation = 180), right (azimuth = 72, elevation = 0) and left (azimuth = 252, elevation = 0). As with Experiment 1, Experiment 2 was split into 4 blocks. All conditions were interleaved pseudo-randomly within each block, with five repeats for each target location and number of sources, for a total of 250 trials.

#### 2.3.3 Experiment 3 (EEG)

In Experiment 3 we used electroencephalography (EEG) to study automatic change detection in the spatialized scene. Participants were exposed to the same stimuli employed in Experiments 1 and 2. However, in this iteration, participants passively listened to the scenes while neural activity was recorded via EEG using a Biosemi Active Two AD-box 703 ADC-17 with 64-channel cap (Biosemi, Netherlands). Participants engaged in a decoy visual task, in which they were asked to press a button when the color of a fixation cross changed from white to green. The cross changed color in 25% of the trials and participants performed at ceiling (mean hit rate: 95.0%, standard deviation: 8.7%) No information on the acoustic scenes was given. Auditory scenes were 7 s in length and composed of five background sources. Appearing sources were added pseudo-randomly in half of the trials, 4 s after trial onset, from one of the five locations described in Experiment 2, and persisted for 3 s. An inter-trial-interval between 2 and 3 s was employed. Each participant was presented with a total of 300 trials, resulting in 30 trials per-appearing-source-location and 150 no-change trials.

#### 2.3.4 Experiment 4 (EEG)

Experiment 4 replicated Experiment 3, but here the stimuli were presented via an 8-loudspeaker array (see Figure 2.2C-D) in a sound booth. This was to test whether we would observe the same neural correlates with a reduced number of loudspeakers. Phantom sources (Günther and Georg, 1976) were used to present sources from all 31 locations, as in Experiments 1, 2 and 3 (see Figure 2.2A-B), with just 8 loudspeakers. Using the IRCAM Spat package (Carpentier et al., 2015) for Max/MSP, this was achieved using amplitude panning that involved the loudspeakers neighboring the desired position of the phantom sources (Ville, 1997). Pilot experiments confirmed that rendering yielded sources were clearly perceived in the intended locations.

### 2.4 Analysis

#### 2.4.1 Behavioral Analysis (Experiments 1 & 2)

To calculate the high-frequency hearing thresholds, the mean of the five frequencies tested (2, 4, 8, 12 and 16kHz) and both ears were taken for each participant (threshold mean: 21.4 dBA, standard deviation: 12.9 dBA). In the change detection task, each participant’s behavioral data, performance and reaction time, were logged during the experiment and extracted offline with custom Python scripts. A “hit” was classed as a button-press at any point after the onset of the appearing source, with the hit rate as the number of trials over the number of appearing source trials. A false alarm was classed as a button press at any point before the onset of the appearing source and on trials in which there was no appearing source. As participants could false alarm on all trials, the false alarm rate was calculated as the number of false alarms over the total number of trials. The criterion was also calculated as a measure of bias on a participant’s likelihood of pressing the button. With the Stanislaw and Todorov (1999) correction on hit rate and false alarm rate, a d’ was calculated as a measure of performance by the z-transform of the hit rate minus the z-transform of the false alarm rate.

Statistical analysis of (1) the effects of the number of concurrent sources and target location or (2) whether the scene was spatialized or not, were performed using general or generalized linear models fitted with *glm* of the Python *statsmodels* package. The details of each model are outlined alongside the relevant results, with the analysis of behavioral performance (correct vs. incorrect responses) modelled via a logistic regression in which the generalized linear model used a binomial distribution. Analysis of other metrics such as reaction time and criterion were modelled using a general linear model. For each model we report the magnitude of coefficients (estimate) of fixed effects of interest, the test statistic for comparison of the coefficients to 0 (*Z*) and its respective p-value (*p*).

#### 2.4.2 EEG Analysis (Experiments 3 & 4)

##### 2.4.2.1 Preprocessing

The raw EEG data underwent processing using the FieldTrip toolbox (Oostenveld et al., 2011). Epochs were extracted from 200 ms before to 700 ms after the change. In “no change” trials, epochs of identical duration extracted around 4 seconds post onset (matching the time at which changes occurred in change trials). Trials were rejected manually based on visual inspection via the FieldTrip user interface, to exclude from the analysis any trials that had a z-score exceeding three, that deviated from the observed trend line, or that displayed variances that were visually larger than neighboring trials. A bandpass filter (two-pass, fifth-order Butterworth) was then applied below 1 Hz and above 30 Hz. To optimize computational efficiency, the data were down sampled to 256 Hz and subsequently de-meaned, subjected to linear detrending, and re-referenced to the average of all channels using FieldTrip’s built-in functions.

To further refine the data for downstream analysis, a second round of manual trial removal was conducted. Channels identified as problematic were removed last, based on visual inspection of variance to identify outliers, resulting in minimal necessity for channel removal. The data then underwent denoising using the "Dynamic Separation of Sources" (DSS) algorithm (de Cheveigné and Simon, 2008), applied across all conditions. The DSS algorithm enhances reproducibility of stimulus-evoked responses across trials (de Cheveigné and Parra, 2014; de Cheveigné and Simon, 2008). The top five components derived per-participant from all trials were retained for further analysis and projected back into sensory space. Trials were averaged and de-meaned per condition, with the process iteratively applied to each group of change/no-change conditions per-location and all change/no-change conditions regardless of location. To address variations in trial numbers in individual change conditions, a comparable number of no-change trials were randomly selected for accurate comparison across conditions.

##### 2.4.2.2 Population analysis

To quantify the response across channels we used a measure of instantaneous power. The root mean square (RMS) value of the average time series from eight channels with the highest amplitudes from the group-averaged 300 ms peak topographic plots (discussed below) was computed for each participant and each pair of conditions. Group-averaged RMS values were then calculated per-condition. These underwent a bootstrap analysis (Tibshirani and Efron, 1993) using 1000 samples per condition. Significance bars were plotted based on a bootstrap resampling approach, with a *p* value threshold set at 0.01 and requiring at least 31 ms of consecutive significant samples.

Topographical analyses were performed to investigate differences between experimental conditions in the specified peak time window. To standardize the number of channels for each participant after pre-processing for group topography analysis, channel interpolation was performed per-participant based on a neighbors-derived weighted average, with an average of seven channels interpolated per-participant. The resultant interpolated data were then group-averaged for each condition, with a 100 ms window around the peak at 0.3 s plotted per-condition. A permutation analysis was employed to assess the statistical significance between pairs of topographies at the peak of interest. To elucidate any relationships between experimental conditions and the observed peaks at 100 ms, 300 ms, and 600 ms, we conducted Spearman’s rank correlation analyses on the interpolated group-averaged topographical data. For each time window and condition, pairwise Spearman’s rank correlations were computed between all condition pairs with false discovery rate correction for multiple comparisons. Results were organized into correlation matrices for each time window, and ultimately revealed no statistically significant correlations between any condition pair.

To determine the optimal sample size for detecting significant differences between change and no-change conditions, a permutation-based analysis was conducted using the group-averaged RMS data. Subset sizes were examined ranging from 5 to 30 participants in increments of 5, performing 100 iterations for each subset size. For each iteration, participants were randomly selected and 1000 permutations were conducted to generate a null distribution. P-values were calculated by comparing observed differences to this null distribution, with a significance threshold of p < 0.01. The stability of p-value patterns across different sample sizes were then compared to those obtained from the full dataset (N = 32).

## 3. RESULTS

### 3.1 Experiment 1: spatializing sources benefits change detection

A trial-based logistic regression was performed to identify differences in performance due to spatialization, as a function of the number of sources and the participants’ high-frequency hearing thresholds. We observed significant main effects of spatialization (β = -3.611, *p* = 0.001; reference = spatialized), number of sources (β = -0.466, *p* < 0.001) and hearing threshold (β = - 0.0522, *p* = 0.036) as well as significant interactions between these predictors (see Table B in supplementary materials). Therefore, a trial-based logistic regression was performed on each condition (spatialized and non-spatialized) separately. Main effects of the number of sources (β = -0.466, *p* < 0.001) and high-frequency hearing threshold (β = -0.0522, *p* = 0.036) were identified in the spatialized condition but not in the non-spatialized condition (see Figure 3.1, or see Table 10.1 to Table 10.3 for the full models in the supplementary materials). For reaction time, a trial-based linear regression revealed no significant main effects. Overall, participants showed better performance for spatialized scenes. Additionally, in spatialized scenes, but not non-spatialized scenes, performance decreased with a growing number of scene sources, and with worse high frequency hearing.

**Figure 3.1.**
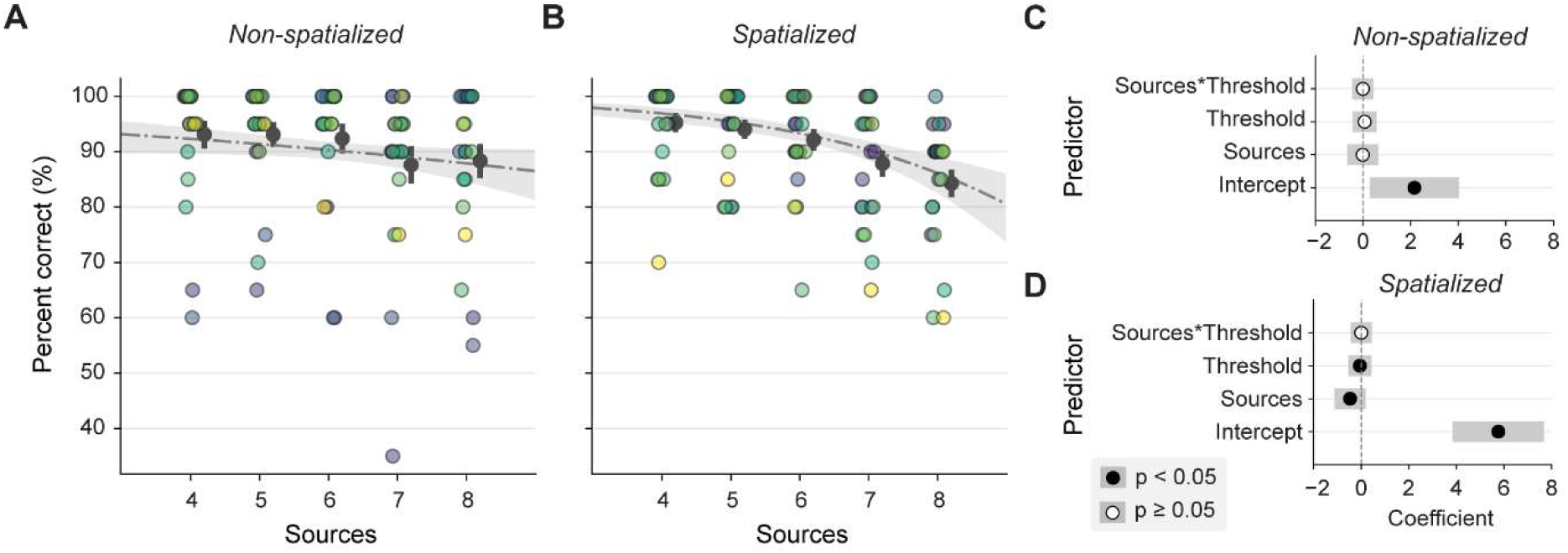
Behavioral performance for change detection in non-spatialized vs. spatialized acoustic scenes. Plots of the performance in each condition (A: Non spatialized, B: Spatialized) with predicted values from the logistic regression in grey. Each data point is colored by the individual participant and the error bars show the mean and standard error across participants. (C-D) Model coefficients for the trial-based logistic regression are displayed for the non-spatialized and spatialized conditions and are colored by whether the predictor showed a significant effect.

### 3.2 Experiment 2: appearing-source’s location modulates performance, dependent on scene complexity

Whereas Experiment 1 investigated performance as a function of increasing background source numbers in non-spatialized versus spatialized scenes, Experiment 2 aimed to assess the impact of the appearing source’s location within spatialized scenes. These scenes were similarly composed of different numbers of background sources. As a Shapiro-Wilk test revealed the data to be non-normal, a participant-based Friedman analysis of variance was performed. This revealed no significant effect of location on the proportion of correct trials (χ2(4) = 4.76, *p* = 0.313), the measure of d’ (χ2(4) = 6.99, *p* = 0.136), or the criterion measure (χ2(4) = 1.26, *p* = 0.869; see Figure 3.2A-C). A trial-based logistic regression similarly identified no main effect of location, but did identify a main effect of the number of sources, where increasing source number decreased the proportion of correct responses (β = -0.394, *p* < 0.001).

**Figure 3.2.**
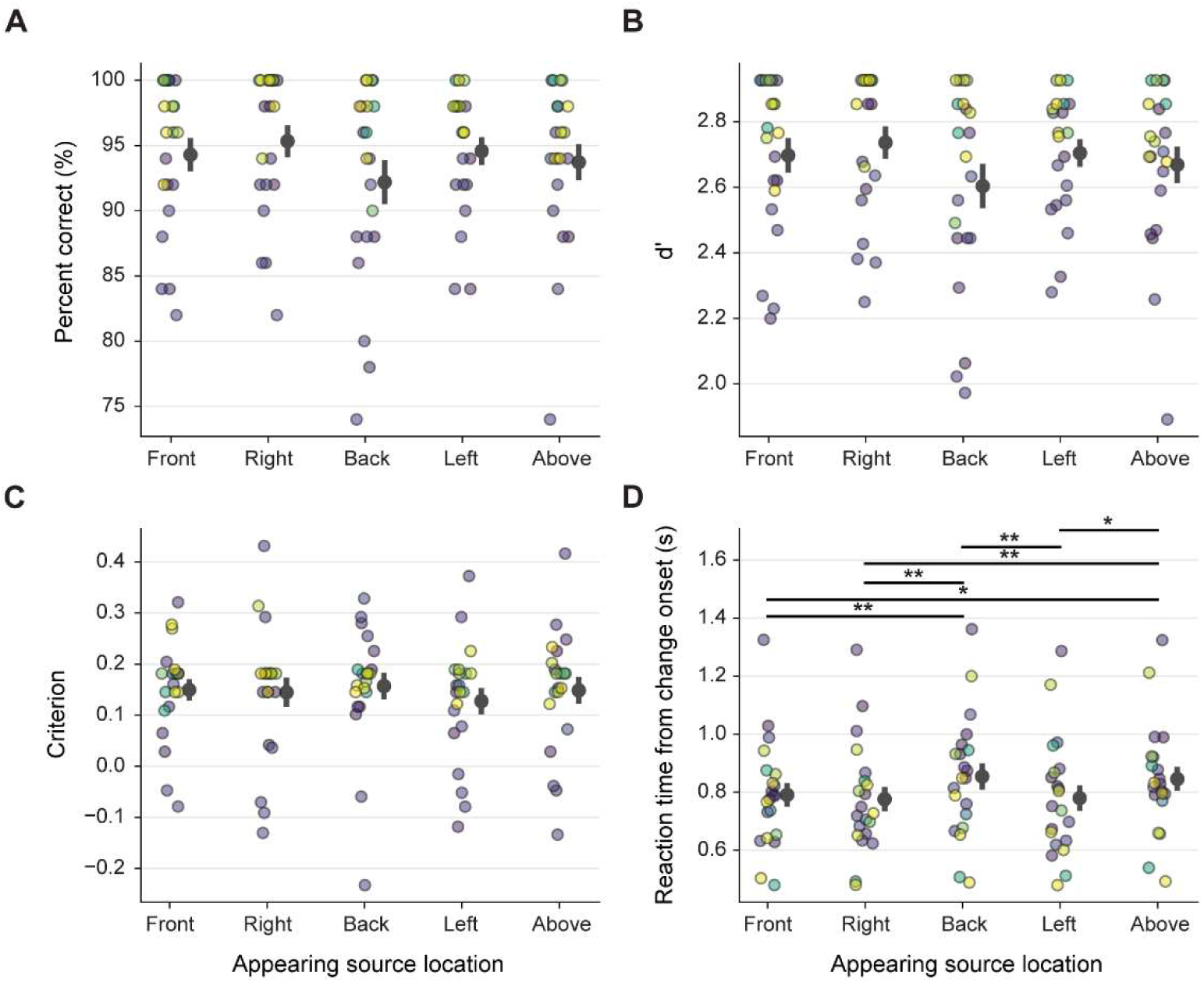
Reaction time is influenced by the location of the appearing source. Performance for each target location as a measure of (A) percent correct trials, (B) d’, (C) criterion and (D) reaction time of participants to the appearing sound source in ‘hit’ trials, all collapsed across the number of sources. Each data point is colored by the individual participant and the error bars show the mean and standard error across participants. ** = p < 0.01, * = p < 0.05.

To assess the effect of appearing source location on reaction time, we collapsed all “hits” across background source conditions. Reaction time was also found to be non-normal (Shapiro-Wilks) and a participant-based Friedman analysis of variance revealed statistically significant differences in reaction time between at least two location conditions (χ2(4) = 25.79, *p* < 0.001, see Figure 3.2D). Post-hoc pairwise comparisons showed significant differences between mostly the back and above locations from the front, left and right locations (Wilcoxon test with holm correction; see Table 10.4 in supplementary for the comparison table).

To further assess the effect of source location and scene complexity on reaction time, a linear regression was performed on each appearing source location condition, split by the number of background sources. This analysis showed a significant interaction term between the above location and the number of sources (*p* = 0.024). Therefore, a trial-based linear regression was performed for each appearing source location separately. Only a significant effect of sources was identified for the above and back appearing sound source locations, where reaction time increases with an increasing number of sources (above: β = 0.0573, *p* < 0.001; back: β = 0.0338, *p* = 0.012; see Figure 3.3).

**Figure 3.3.**
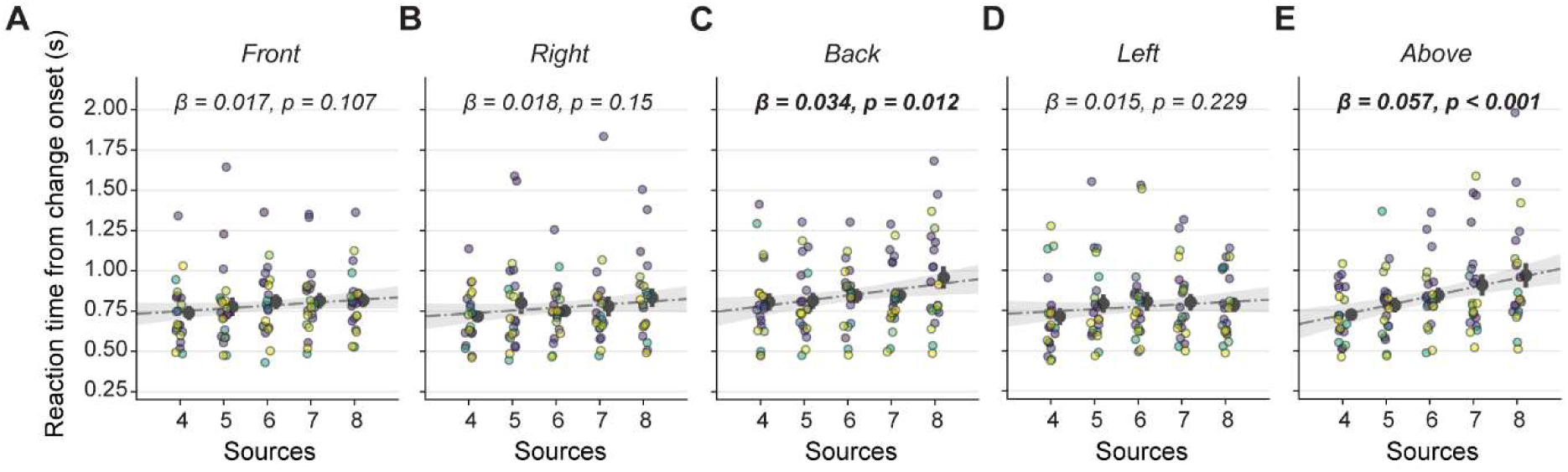
Reaction time plotted for each participant for each appearing source location. (A) front, (B) right, (C) back, (D) left, (E) above and the number of sources present for ‘hit’ trials. The confidence intervals and predicted mean values from the linear regression for each target location is shown with a gray shaded area and dashed line, respectively. The coefficient and p value for the effect of ‘sources’ on reaction time is annotated on each subplot.

In Experiment 1, spatializing sound sources provided a significant performance benefit for detecting new appearing sources compared to non-spatialized presentation. However, this benefit declined as the number of background sources increased and was modulated by participants’ high-frequency hearing thresholds, with poorer thresholds leading to worse performance in spatialized conditions. In Experiment 2, while the proportion of correctly detected appearing sources did not differ as a function of appearing source location, reaction times were significantly slower when sources appeared from above or behind the listener compared to frontal or lateral locations. This location-based reaction time effect was exacerbated as the number of background sources increased, particularly for sources appearing from above.

### 3.3 Experiment 3: neural correlates of change detection are independent of change location

In Experiments 1 and 2, participants were given an active listening task, with sources added to the scene from different locations. In Experiment 3, we investigated neural correlates of passive change detection; participants were instead given a visual decoy task while passively exposed to similar scenes and appearing sources/locations, while their neural activity was recorded via EEG. Compared with the group-averaged baseline-corrected time courses in which no appearing source was added (the “no change” condition), group-averaged appearing source time courses, averaged across locations (the “change” condition), showed a significant change complex between 200-500 ms post-change-onset (see Figure 3.4A). The statistical analysis uncovered significant differences between “change” and “no change” conditions (*p* < 0.01) with peaks emerging around 200 ms and 300 ms, and a third late peak around 600 ms. When separating the time courses by appearing source location (see Figure 3.4B-F), no significant differences were observed between location conditions, but each change location was significantly different from the “no change” condition (derived from a subset of “no change” trials for equal comparison).

**Figure 3.4.**
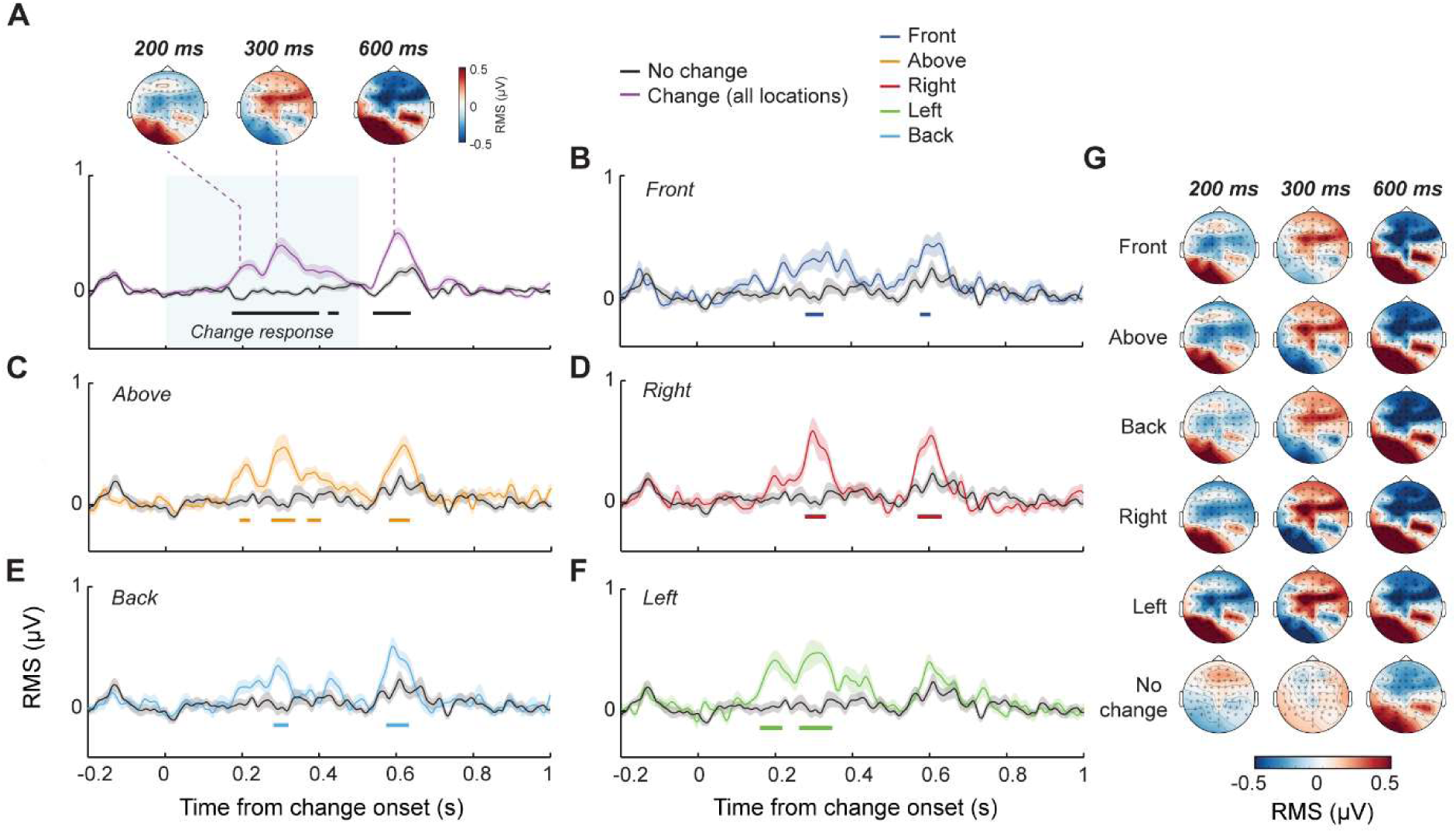
Neural correlates of change detection in Experiment 3. Baseline-corrected RMS time series (see methods), with significance bars between condition comparison (p < 0.01). A). Group-averaged across all “appearing source” and all “no change” conditions; inset topographies elicited by each peak. B-F). Group-averaged across individual conditions and a matched subset of “no change” trials. (G) Group Topographies separated by appearing source location conditions; 100 ms window surrounding the peak at 200 ms, 300 ms and 600 ms from change onset.

Group scalp topographies were plotted after the per-participant channel interpolation was performed, with topographic maps plotted per condition based on a 100 ms window surrounding the peaks at 200 ms, 300 ms, and 600 ms (see Figure 3.4B-G). A post-hoc permutation analysis was then performed to further assess location-based differences in neural activity patterns through a pairwise comparison of each condition. Ultimately, no statistically significant differences between topographic features were detected across appearing source condition pairs at the specified peak time, with a significance threshold set at *p* < 0.05, suggestive of a shared cortical network in the processing of unattended spatial changes within the confines of our stimuli and experimental design. An analysis between sample size and the similarity of *p* values to those obtained from the full population was also conducted. The *p* value patterns for N = 20 participants closely resembled those for N = 32 participants (*p* = 0.023), suggesting diminishing returns for sample sizes beyond 20.

### 3.4 Experiment 4: Neural correlates of changes detection persist with a spatial scene of phantom sources

In Experiment 4, we examined neural activity under conditions similar to those in Experiment 3, with the modification of using phantom sources in a reduced speaker setup. A comparison of group-averaged, baseline-corrected time courses revealed that, relative to the "no change" condition, the "change" condition exhibited a significant change complex between 150-320 ms post-change-onset (*p* < 0.01; see Figure 3.5A). This complex was characterized by two peaks of activity, emerging around 210 ms and 290 ms. Additionally, the analysis identified a second phase of activity occurring between approximately 400-600 ms, marked by a single peak around 550 ms post-change-onset. As in Experiment 3, no significant differences were observed when separating the RMS time courses by the location of the appearing source. However, each change location exhibited a similar two-phase response pattern compared to a randomly drawn, matching subset of "no change" trials (see Figure 3.5, B-F). The group scalp topographies were analyzed for peaks at 210 ms, 294 ms, and 554 ms, using the same interpolation procedure and window lengths as in Experiment 3. Consistent with prior findings, a post-hoc permutation analysis detected no significant differences in the topographic features of change-evoked neural activity across appearing source conditions for each of these peaks.

**Figure 3.5.**
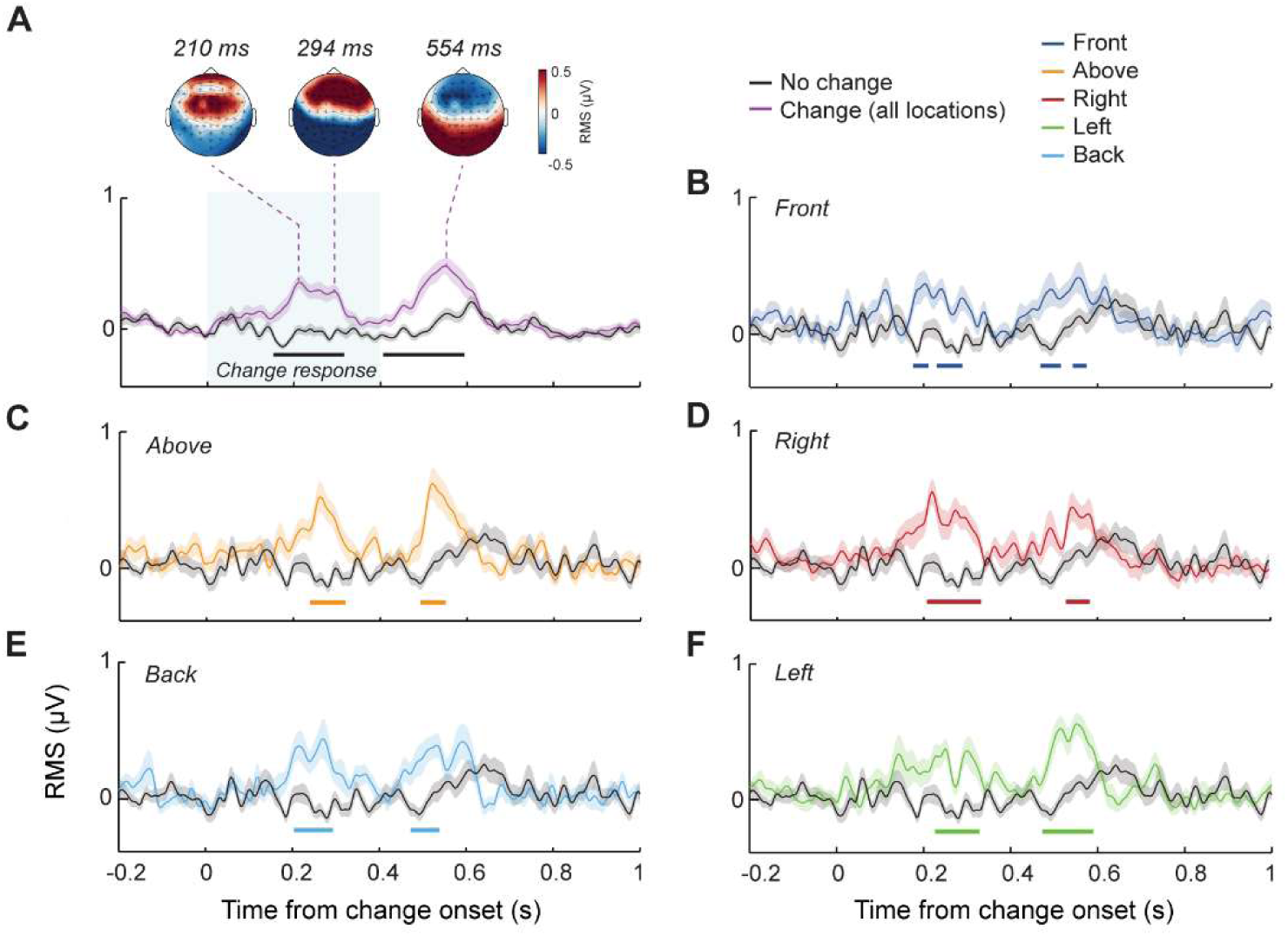
Neural correlates of change detection in Experiment 4. Baseline-corrected RMS time series, with significance bars between condition comparison (p < 0.01). A). Group-averaged across all “appearing source” and all “no change” conditions; inset topographies elicited by each peak. B-F). Group-averaged across individual conditions and a matched subset of “no change” trials.

To confirm the robustness of the latency of the neural responses and a lack of difference between spatial location conditions, we aggregated the EEG data across experiments 3 and 4. The only difference between Experiment 3 and 4 was the loudspeaker arrangement; Experiment 3 used 31 loudspeakers and Experiment 4 used 8 loudspeakers but with 31 phantom sources. These results are provided in Figure 3.6. Consistent with the results from each of the experiments individually, no significant differences were observed between the different change location conditions.

**Figure 3.6.**
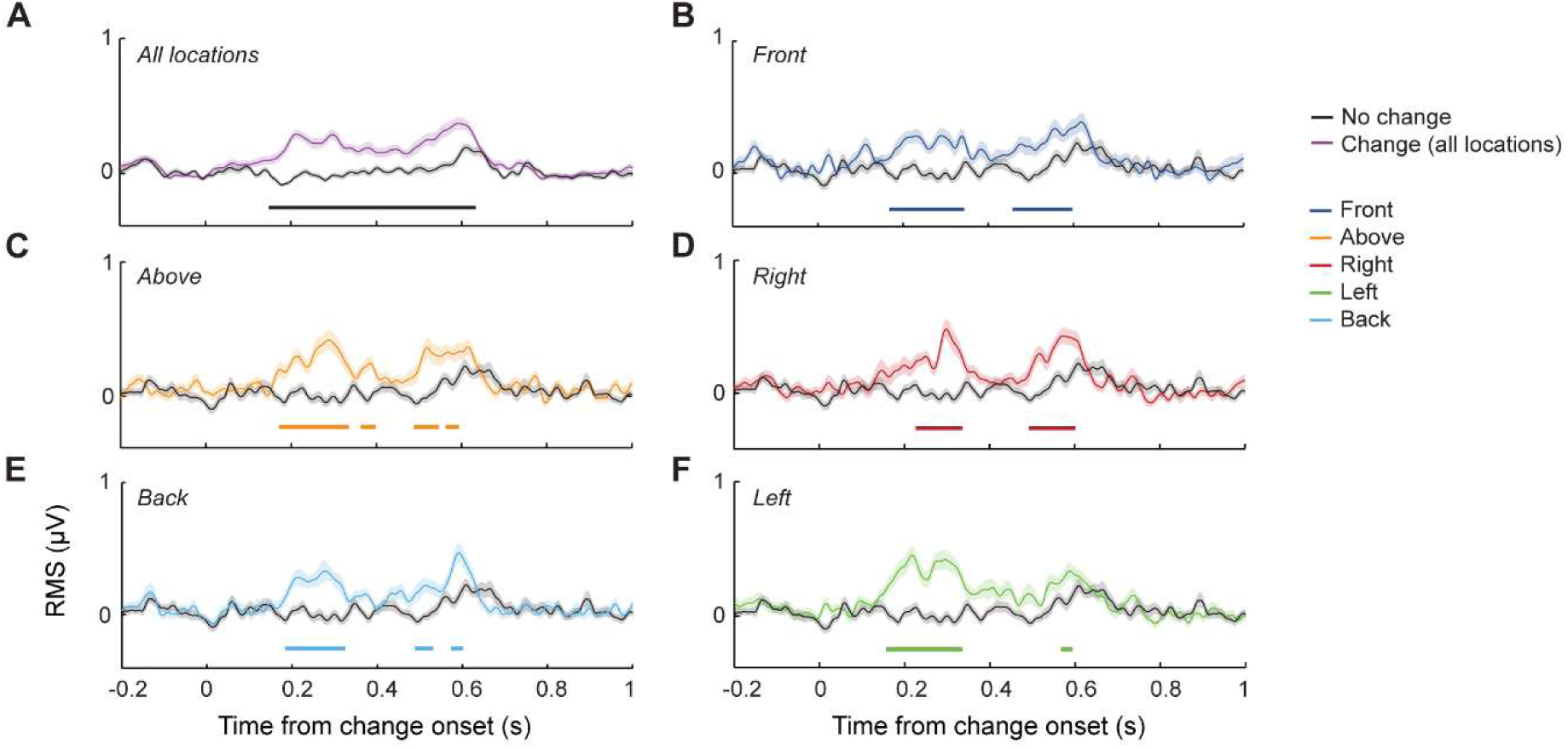
Neural correlates of change detection aggregated across Experiment 3 and 4. Baseline-corrected RMS time series, with significance bars between condition comparison (p < 0.01). A). Group-averaged across all “appearing source” and all “no change” conditions. B-F). Group-averaged across individual conditions and a matched subset of “no change” trials.

## 4. DISCUSSION

To investigate listeners’ sensitivity to changes in complex, spatial auditory environments, we created naturalistic “chimeric” sounds designed to mimic the spectrotemporal complexity of real-world environments while minimizing semantic bias (Vanden Bosch der Nederlanden et al., 2020). These sounds were largely broadband, thus supporting localization in both azimuth and elevation through binaural (interaural level and timing differences) and monaural (spectral) auditory cues. The multi-loudspeaker spherical array allowed us to position sound sources around and above the listener. In Experiments 1 and 2, participants tracked auditory scenes for changes, with performance influenced by the number, spatial arrangement, and location of sound sources, specifically when sounds emerged from behind or above the listener. In Experiments 3 (using the multi-loudspeaker array) and 4 (using a reduced speaker array implementing phantom sources), we investigated automatic change detection with EEG revealing robust neural correlate of change detection.

### 4.1 Spatial separation of sound sources impacts auditory change detection

Most prior research on auditory change detection has examined non-spatialized audio (de Kerangal et al., 2021; Gregg et al., 2014; Gregg and Snyder, 2012; Sohoglu and Chait, 2016a) or have spatialized the auditory streams but did not examine its impact (Puschmann et al., 2013). Several studies have examined the influence of spatialization, yielding mixed findings: some report enhanced change detection when sources are spatially separated (Aman et al., 2021; Eramudugolla et al., 2005; Gaston et al., 2017), while others note impairments (Gregg and Samuel, 2008). In our study, Experiment 1 revealed that spatially separating sound sources facilitated behavioral change detection compared to conditions where sources were non-spatialized. This finding aligns with previous studies using both free-field (Aman et al., 2021; Gaston et al., 2017) and virtual spatialized sounds (Eramudugolla et al., 2005), both reporting an impairment in detection performance when sound sources were co-located. Notably, we also observed that reduced high-frequency hearing thresholds led to lower change detection accuracy, but only in the spatialized condition. Since high-frequency monaural spectral cues play an important role in spatial hearing (Best et al., 2005; Blauert, 1997), these results suggest that high-frequency hearing acuity is a significant factor in effective change detection within spatialized auditory scenes. As a whole, these findings are consistent with well-documented effects of spatial release from masking in auditory scene analysis (Bregman, 1994; Kidd et al., 1998; Shinn-Cunningham, 2008; Snyder and Alain, 2007), and demonstrate that spatial release from masking improves not only our perception of attended stimuli, but also enhances listeners’ abilities to discern changes in complex acoustic scenes.

Prior research consistently demonstrates that increasing the number of auditory streams impairs change detection performance (Constantino et al., 2012; Petsas et al., 2016; Sohoglu and Chait, 2016a). This pattern was evident in our spatialized condition, where participants’ performance declined as the number of concurrent sound sources increased. In contrast, the non-spatialized condition showed no such effect, with the number of sources having no measurable impact on change detection. This suggests that different processing mechanisms may have been invoked to support change detection for these two conditions. In the non-spatialized condition, the absence of spatial cues for source segregation meant that adding more auditory streams may not have increased the perceptual complexity of the scene. Listeners might instead have relied mainly on simple amplitude changes to detect the appearance of a new sound source. Conversely, in the spatialized condition, spatialized sources were more readily perceived as distinct auditory streams, and the addition of more sources may therefore have increased the perceptual complexity of the scene. Given this, listeners are able to direct their attention towards individual streams, thus potentially shifting focus away from where new sources might appear (Alain and Arnott, 2000; Best et al., 2006; Kishline et al., 2013; Murphy et al., 2017). This could partially explain the reduced change detection performance in spatialized scenes with a high number of concurrent sources. Although the underlying reasons for differences between spatialized and non-spatialized conditions remain unclear, it is evident that spatial separation of sound sources significantly impacts auditory change detection. Future research on auditory change detection should consider this effect when incorporating spatialized stimuli.

### 4.2 Slower reaction times for sources that appear above or behind the listener

In Experiment 2, we examined how the spatial location of the appearing sound source influenced change detection by comparing five location conditions: left, right, front, above and behind the listener. Prior studies have demonstrated that the location of an auditory target in quiet impacts behavior, with faster reaction times in response to sounds that originate from locations outside the visual field (Asutay and Västfjäll, 2015; Drossos et al., 2015; Tajadura-Jiménez et al., 2010a, 2010b). This “rear-bias” has been shown only for naturalistic auditory stimuli with semantic content, and not for simple tonal stimuli (Olszanowski et al., 2023). To our knowledge though, no previous work has examined how the location of an auditory target affects behavior when it appears in a spatialized scene, rather than in quiet. In our study, which used concurrent non-semantic sounds, we found that reaction times increased when changes originated from either above or behind the listener, and this was exacerbated as the number of sources increased.

One possible interpretation of this location-specific delay in reaction time is that it arises from the interaction between the acoustic waves and anatomic elements of the head, torso and pinnae. These may subtly favor sounds originating from the front by attenuating those from above or behind (Carlile and Pralong, 1994), but to different degrees depending on the individual listeners. However, considering the complexity of the auditory scenes in our experiments – which included background sounds behind the listener and variable sound envelopes - it seems unlikely that such filtering effects alone would consistently and substantially increase reaction times. Furthermore, if these acoustic variations were substantial, we would expect corresponding patterns in the underlying neural activity, with delayed or reduced responses for sources appearing behind the listener. No such differences were observed in the EEG data, although some caution is necessary when interpreting a null result.

An alternative interpretation of this delayed reaction time is that it arises from a spatial-attention bias that favors sounds originating from the direction of the listener’s gaze, which in our experiments was oriented towards the front. Previous research has shown that reaction times are fastest when a listener’s gaze and spatial attention are aligned (Pomper and Chait, 2017). This gaze focus of attention may lead to “change-deafness” or a delay in change detection for sound sources appearing outside the (visually-) attended region (Demany et al., 2017), while enhancing detection for changes in attended areas (Best et al., 2023; Eramudugolla et al., 2005; Irsik et al., 2016). This spatial-attention account is consistent with the finding that the most pronounced delays in reaction time occurred in the condition with the highest number of sources, where attentional load would be greatest. Thus, rather than attributing these reaction time differences between locations to acoustic filtering, the interaction between attentional mechanisms and location may provide a more comprehensive explanation.

### 4.3 Change events evoke robust neural responses

Our EEG data from Experiments 3 and 4 provided the first evidence that robust neural correlates of auditory change detection can be identified within complex, naturalistic, spatialized acoustic scenes under passive listening conditions. In these conditions, change detection responses emerged approximately 200 ms following the onset of a change; about 100 ms later than that observed in studies that used simpler, non-spatialized stimuli (Sohoglu and Chait, 2016a). Previous research on simple auditory scenes has shown that the detection of an appearing sound source relies on sensitivity to spectral transients – specifically an increase in energy in a previously silent frequency (Constantino et al., 2013). Brain imaging studies link spectral transient responses to abrupt and early neural markers, such as the M50 response, a peak in the neural response occurring around 70 ms after sound onset (Sohoglu and Chait, 2016a). The neural responses that we observed in our data may lack this early response due to the absence of pronounced transients in the complex spectra of our stimuli. Alternately, our complex stimuli may have produced transients at different time points in each trial depending on the interaction between the newly-appearing source and the background sources, and therefore the early response was absent in the average.

Prior studies on the neural correlates of change detection noted additional peaks in the neural responses that were associated with the perceptual “pop-out” of new sounds and their subsequent attentional tracking (Gregg and Snyder, 2012; Puschmann et al., 2013; Sohoglu and Chait, 2016a). These later responses have been correlated with behaviorally-relevant processes that contribute to change detection outcomes (Gregg and Snyder, 2012; Puschmann et al., 2013) but occur regardless of whether changes are consciously detected (Sohoglu and Chait, 2016a). Our observed 200 ms peaks likely correspond to these later stages of source tracking. These responses consistently appeared in both Experiments 3 and 4, underscoring their robustness. Topographical patterns near these activity peaks suggest ERP-like components resembling the MMN, N2bm and P3b complexes that have been reported in previous research (Puschmann et al., 2013). These peaks may reflect the sequential processing of auditory stimuli, encompassing the initial detection of a new sound stream, attentional capture, and the shift from detection to higher-order cognitive processing (Folstein and Van Petten, 2008; Näätänen et al., 2007; Polich, 2007).

We also detected an additional peak at 600 ms which has not been reported in previous work. The significance of this peak remains speculative. It is unclear whether this peak represents an additional correlate of source tracking or if it is a stimulus-specific response linked to the temporally varied envelope of the stimuli that we used. Further research, employing a different set of chimeric sounds, would be helpful to understand this peak’s origin.

### 4.4 Change-evoked neural responses are independent of location

Though the behavioral results indicated delayed reaction times for sound sources appearing above and behind the listener, no such differences between locations were observed in the EEG responses. Change-evoked responses across all studied locations showed comparable amplitude and latency, thereby suggesting equal salience. Even when combining data from both EEG experiments (N = 62), no differences between locations emerged, suggesting that the null effect of location is unlikely to be due to a lack of power. Thus, while null results must always be interpreted with caution, our findings support the conclusion that, by 200 ms post-onset, scene changes are processed with similar latencies regardless of spatial location. These findings imply that the behavioral effects of location may arise from later attentional or decision-making processes, which are not reflected in the early sensory components measured by EEG. This explanation echoes the modulatory effects observed on evoked responses to spatialized speech-in-noise location changes compared to speech-in-silence in previous research (Ozmeral & Menon, 2023). Alternately, the six concurrent sources used in Experiments 3 and 4 may have been insufficient to elicit location-dependent effects. Reaction time impairments for changes above and behind the listener were most pronounced with eight concurrent sources. Thus, the lower number of sources in Experiments 3 and 4 may have diminished location-based differences in neural responses.

The absence of observed neural differences between appearing source locations does not rule out their existence. Such effects may be subtle and masked by EEG’s susceptibility to noise, further complicated by the complexity of the stimuli. Location-specific effects could still emerge with alternative paradigms or more advanced analysis. Future research on the neural correlates of change detection in complex auditory scenes could benefit from increasing scene complexity to elevate perceptual load. Additionally, studies requiring listeners to actively attend to the auditory scene may reveal spatial differences in neural responses that were not apparent under passive listening conditions. Several studies have shown that attention can modulate neural signals (Ozmeral et al., 2021), suggesting that an active listening task might uncover effects related to the location of change detection. The role of attention in facilitating change detection in such scenes remains a topic ripe for investigation, as the cognitive processes underlying neural mechanisms, and their relation to auditory attentional deficits remain an important field of inquiry (Razzaghipour et al., 2024).

### 4.5 Conclusion

We sought to contribute to the available toolkit for investigating auditory spatial processing and change detection. Our findings reveal the relationship between spatial hearing, spectrotemporal complexity, and attention in auditory change detection. By simulating naturalistic auditory conditions whilst minimizing semantic information, this study advances our understanding of the neurocognitive mechanisms supporting change detection in complex acoustic scenes. Behavioral data show that spatial cues, in combination with the complexity of the scene and hearing ability, modulate change detection responses, with increasing source numbers reducing responsiveness in spatialized settings. Our EEG findings provide novel insights into the cortical processing of unattended spatial auditory change detection and overall, our data underscore the role of attention and spatial acuity in shaping behavioral and neural responses. These insights stress the need to consider attentional effects in experimental designs for complex spatialized acoustic scenes.

Future studies should leverage task-based auditory paradigms to determine whether spatial cues affect later stages of auditory processing, potentially influencing decision-making in complex acoustic scenes. Our focus on appearance-based change detection, as opposed to disappearances, may have highlighted unique mechanisms. Given our use of stimuli with minimal semantic associations, future work could explore the impact of semantic features on attentional modulation in spatialized scenes, providing further insight into reaction times and task accuracy in dynamic auditory environments. Finally, paradigms such as ours, that involve passive spatialized change detection, hold promise in fields such as hearing aid signal processing, where current algorithms may limit users’ ability to accurately localize sounds. This approach may offer a way to assess spatial sound virtualization algorithms used in augmented and virtual reality, providing empirical measures of their effectiveness based on neural responses.

## ACKNOWLEDGEMENTS

This study was funded by the William Demant Foundation (Case no. 21-2519). We thank Zyque So for assistance with data collection.

## 5. AUTHOR CONTRIBUTIONS

**Katarina C. Poole:** Methodology, Software, Formal analysis, Investigation, Data Curation, Writing – Original Draft, Writing – Review & Editing, Visualization. **Drew Cappotto:** Methodology, Formal analysis, Investigation, Writing – Original Draft. **Vincent Martin:** Methodology, Software, Formal analysis, Investigation, Writing – Review & Editing. **Jakub Sztandera:** Validation, Formal analysis, Investigation. **Maria Chait:** Methodology, Conceptualization, Software, Writing – Review & Editing, Supervision, Funding acquisition. **Lorenzo Picinali:** Conceptualization, Writing – Review & Editing, Supervision, Project administration, Funding acquisition. **Martha Shiell:** Conceptualization, Writing – Review & Editing, Supervision, Funding acquisition.

## REFERENCES

Alain, C., Arnott, S.R., 2000. Selectively attending to auditory objects. FBL 5, 202–212. 10.2741/alain

Aman, L., Picken, S., Andreou, L.-V., Chait, M., 2021. Sensitivity to temporal structure facilitates perceptual analysis of complex auditory scenes. Hearing Research 400, 108111. 10.1016/j.heares.2020.108111

Asutay, E., Västfjäll, D., 2015. Attentional and emotional prioritization of the sounds occurring outside the visual field. Emotion 15, 281–286. 10.1037/emo0000045

Best, V., Boyd, A.D., Sen, K., 2023. An Effect of Gaze Direction in Cocktail Party Listening. Trends in Hearing 27, 23312165231152356. 10.1177/23312165231152356

Best, V., Carlile, S., Jin, C., van Schaik, A., 2005. The role of high frequencies in speech localization. The Journal of the Acoustical Society of America 118, 353–363. 10.1121/1.1926107

Best, V., Gallun, F.J., Ihlefeld, A., Shinn-Cunningham, B.G., 2006. The influence of spatial separation on divided listening. The Journal of the Acoustical Society of America 120, 1506–1516. 10.1121/1.2234849

Blauert, J., 1997. Spatial Hearing: The Psychophysics of Human Sound Localization. MIT Press.

Bregman, A.S., 1994. Auditory scene analysis: The perceptual organization of sound. MIT press.

Carlile, S., Pralong, D., 1994. The location-dependent nature of perceptually salient features of the human head-related transfer functions. The Journal of the Acoustical Society of America 95, 3445–3459. 10.1121/1.409965

Carpentier, T., Noisternig, M., Warusfel, O., 2015. Twenty Years of Ircam Spat: Looking Back, Looking Forward, in: 41st International Computer Music Conference (ICMC). Denton, TX, United States, pp. 270–277.

Constantino, F.C., Pinggera, L., Chait, M., 2013. The Role of Sensitivity to Transients in the Detection of Appearing and Disappearing Objects in Complex Acoustic Scenes, in: Moore, B.C.J., Patterson, R.D., Winter, I.M., Carlyon, R.P., Gockel, H.E. (Eds.), Basic Aspects of Hearing. Springer, New York, NY, pp. 183–192. 10.1007/978-1-4614-1590-9_21

Constantino, F.C., Pinggera, L., Paranamana, S., Kashino, M., Chait, M., 2012. Detection of Appearing and Disappearing Objects in Complex Acoustic Scenes. PLOS ONE 7, e46167. 10.1371/journal.pone.0046167

de Cheveigné, A., Parra, L.C., 2014. Joint decorrelation, a versatile tool for multichannel data analysis. NeuroImage 98, 487–505. 10.1016/j.neuroimage.2014.05.068

de Cheveigné, A., Simon, J.Z., 2008. Denoising based on spatial filtering. Journal of Neuroscience Methods 171, 331–339. 10.1016/j.jneumeth.2008.03.015

de Kerangal, M., Vickers, D., Chait, M., 2021. The effect of healthy aging on change detection and sensitivity to predictable structure in crowded acoustic scenes. Hearing Research, Stimulus-specific adaptation, MMN and predicting coding 399, 108074. 10.1016/j.heares.2020.108074

Demany, L., Bayle, Y., Puginier, E., Semal, C., 2017. Detecting temporal changes in acoustic scenes: The variable benefit of selective attention. Hearing Research 353, 17–25. 10.1016/j.heares.2017.07.013

Drossos, K., Floros, A., Giannakoulopoulos, A., Kanellopoulos, N., 2015. Investigating the Impact of Sound Angular Position on the Listener Affective State. IEEE Transactions on Affective Computing 6, 27–42. 10.1109/TAFFC.2015.2392768

Eramudugolla, R., Irvine, D.R.F., McAnally, K.I., Martin, R.L., Mattingley, J.B., 2005. Directed Attention Eliminates ‘Change Deafness’ in Complex Auditory Scenes. Current Biology 15, 1108–1113. 10.1016/j.cub.2005.05.051

Folstein, J.R., Van Petten, C., 2008. Influence of cognitive control and mismatch on the N2 component of the ERP: A review. Psychophysiology 45, 152–170. 10.1111/j.1469-8986.2007.00602.x

Gaston, J., Dickerson, K., Hipp, D., Gerhardstein, P., 2017. Change deafness for real spatialized environmental scenes. Cogn. Research 2, 29. 10.1186/s41235-017-0066-3

Gregg, M.K., Irsik, V.C., Snyder, J.S., 2014. Change deafness and object encoding with recognizable and unrecognizable sounds. Neuropsychologia 61, 19–30. 10.1016/j.neuropsychologia.2014.06.007

Gregg, M.K., Samuel, A.G., 2008. Change deafness and the organizational properties of sounds. Journal of Experimental Psychology: Human Perception and Performance 34, 974–991. 10.1037/0096-1523.34.4.974

Gregg, M.K., Snyder, J.S., 2012. Enhanced sensory processing accompanies successful detection of change for real-world sounds. NeuroImage 62, 113–119. 10.1016/j.neuroimage.2012.04.057

Günther, T., Georg, P., 1976. Localization of Lateral Phantom-Sources. Journal of the Audio Engineering Society.

Huron, D., 1989. Voice Denumerability in Polyphonic Music of Homogeneous Timbres. Music Perception 6, 361–382. 10.2307/40285438

Irsik, V.C., Vanden Bosch der Nederlanden, C.M., Snyder, J.S., 2016. Broad attention to multiple individual objects may facilitate change detection with complex auditory scenes. Journal of Experimental Psychology: Human Perception and Performance 42, 1806–1817. 10.1037/xhp0000266

Ju, U., Chuang, L.L., Wallraven, C., 2022. Acoustic Cues Increase Situational Awareness in Accident Situations: A VR Car-Driving Study. IEEE Transactions on Intelligent Transportation Systems 23, 3281–3291. 10.1109/TITS.2020.3035374

Kidd, G., Jr., Mason, C.R., Rohtla, T.L., Deliwala, P.S., 1998. Release from masking due to spatial separation of sources in the identification of nonspeech auditory patterns. The Journal of the Acoustical Society of America 104, 422–431. 10.1121/1.423246

Kishline, L., Larson, E., Maddox, R.K., Lee, A.K., 2013. Selective and divided attention: Spatial and pitch “spotlights” in a non-semantic task. The Journal of the Acoustical Society of America 134, 4230. 10.1121/1.4831543

Murphy, S., Spence, C., Dalton, P., 2017. Auditory perceptual load: A review. Hearing Research, Annual Reviews 2017 352, 40–48. 10.1016/j.heares.2017.02.005

Näätänen, R., Paavilainen, P., Rinne, T., Alho, K., 2007. The mismatch negativity (MMN) in basic research of central auditory processing: A review. Clinical Neurophysiology 118, 2544– 2590. 10.1016/j.clinph.2007.04.026

Olszanowski, M., Frankowska, N., Tołopiło, A., 2023. “Rear bias” in spatial auditory perception: Attentional and affective vigilance to sounds occurring outside the visual field. Psychophysiology 60, e14377. 10.1111/psyp.14377

Oostenveld, R., Fries, P., Maris, E., Schoffelen, J.-M., 2011. FieldTrip: Open Source Software for Advanced Analysis of MEG, EEG, and Invasive Electrophysiological Data. Computational Intelligence and Neuroscience 2011, 156869. 10.1155/2011/156869

Ozmeral, E.J., Eddins, D.A., Eddins, A.C., 2021. Selective auditory attention modulates cortical responses to sound location change in younger and older adults. Journal of Neurophysiology 126, 803–815. 10.1152/jn.00609.2020

Ozmeral, E.J., Menon, K.N., 2023. Selective auditory attention modulates cortical responses to sound location change for speech in quiet and in babble. PLOS ONE 18, e0268932. 10.1371/journal.pone.0268932

Pantev, C., Eulitz, C., Hampson, S., Ross, B., Roberts, L.E., 1996. The Auditory Evoked “Off” Response: Sources and Comparison with the"On" and the “Sustained” Responses. Ear and Hearing 17, 255.

Pavani, F., Turatto, M., 2008. Change perception in complex auditory scenes. Perception & Psychophysics 70, 619–629. 10.3758/PP.70.4.619

Petsas, T., Harrison, J., Kashino, M., Furukawa, S., Chait, M., 2016. The effect of distraction on change detection in crowded acoustic scenes. Hearing Research 341, 179–189. 10.1016/j.heares.2016.08.015

Phillips, D.P., Hall, S.E., Boehnke, S.E., 2002. Central auditory onset responses, and temporal asymmetries in auditory perception. Hearing Research 167, 192–205. 10.1016/S0378-5955(02)00393-3

Polich, J., 2007. Updating P300: An integrative theory of P3a and P3b. Clinical Neurophysiology 118, 2128–2148. 10.1016/j.clinph.2007.04.019

Pomper, U., Chait, M., 2017. The impact of visual gaze direction on auditory object tracking. Sci Rep 7, 4640. 10.1038/s41598-017-04475-1

Pratt, H., Starr, A., Michalewski, H.J., Bleich, N., Mittelman, N., 2008. The auditory P50 component to onset and offset of sound. Clinical Neurophysiology 119, 376–387. 10.1016/j.clinph.2007.10.016

Puschmann, S., Sandmann, P., Ahrens, J., Thorne, J., Weerda, R., Klump, G., Debener, S., Thiel, C.M., 2013. Electrophysiological correlates of auditory change detection and change deafness in complex auditory scenes. NeuroImage 75, 155–164. 10.1016/j.neuroimage.2013.02.037

Salorio-Corbetto, M., Williges, B., Lamping, W., Picinali, L., Vickers, D., 2022. Evaluating Spatial Hearing Using a Dual-Task Approach in a Virtual-Acoustics Environment. Front. Neurosci. 16. 10.3389/fnins.2022.787153

Shinn-Cunningham, B.G., 2008. Object-based auditory and visual attention. Trends in Cognitive Sciences 12, 182–186. 10.1016/j.tics.2008.02.003

Smith, Z.M., Delgutte, B., Oxenham, A.J., 2002. Chimaeric sounds reveal dichotomies in auditory perception. Nature 416, 87–90. 10.1038/416087a

Snyder, J.S., Alain, C., 2007. Toward a neurophysiological theory of auditory stream segregation. Psychological Bulletin 133, 780–799. 10.1037/0033-2909.133.5.780

Snyder, J.S., Yerkes, B.D., Pitts, M.A., 2015. Testing domain-general theories of perceptual awareness with auditory brain responses. Trends in Cognitive Sciences 19, 295–297. 10.1016/j.tics.2015.04.002

Sohoglu, E., Chait, M., 2016a. Neural dynamics of change detection in crowded acoustic scenes. Neuroimage 126, 164–172. 10.1016/j.neuroimage.2015.11.050

Sohoglu, E., Chait, M., 2016b. Detecting and representing predictable structure during auditory scene analysis [WWW Document]. eLife. 10.7554/eLife.19113

Stanislaw, H., Todorov, N., 1999. Calculation of signal detection theory measures. Behavior Research Methods, Instruments, & Computers 31, 137–149. 10.3758/BF03207704

Tajadura-Jiménez, A., Larsson, P., Väljamäe, A., Västfjäll, D., Kleiner, M., 2010a. When room size matters: Acoustic influences on emotional responses to sounds. Emotion 10, 416–422. 10.1037/a0018423

Tajadura-Jiménez, A., Väljamäe, A., Asutay, E., Västfjäll, D., 2010b. Embodied auditory perception: The emotional impact of approaching and receding sound sources. Emotion 10, 216–229. 10.1037/a0018422

Tibshirani, R.J., Efron, B., 1993. An introduction to the bootstrap. Monographs on statistics and applied probability 57, 1–436.

Uhrig, S., Perkis, A., Möller, S., Svensson, U.P., Behne, D.M., 2022. Effects of Spatial Speech Presentation on Listener Response Strategy for Talker-Identification. Front. Neurosci. 15. 10.3389/fnins.2021.730744

Vanden Bosch der Nederlanden, C.M., Zaragoza, C., Rubio-Garcia, A., Clarkson, E., Snyder, J.S., 2020. Change detection in complex auditory scenes is predicted by auditory memory, pitch perception, and years of musical training. Psychological Research 84, 585–601. 10.1007/s00426-018-1072-x

Ville, P., 1997. Virtual Sound Source Positioning Using Vector Base Amplitude Panning. Journal of the Audio Engineering Society 45, 456–466.

