## Supplementary material for "Assessing Behavioral and Neural Correlates of Change Detection in Spatialized Acoustic Scenes"

| | | | | | CI 95% | |
|---|---|---|---|---|---|---|
| Fixed effects | Estimate | Standard error | Z | p value | Lower | Upper |
| **Intercept** | 5.763 | 0.752 | 7.665 | < **0.001** | 4.289 | 7.236 |
| **Sources** | -0.466 | 0.112 | -4.148 | < **0.001** | -0.686 | -0.246 |
| **Spatialization (ref = Spatialized)** | -3.611 | 1.051 | -3.437 | **0.001** | -5.670 | -1.552 |
| **Sources:Spatialization** | 0.450 | 0.159 | 2.832 | **0.005** | 0.138 | 0.761 |
| **Threshold** | -0.052 | 0.025 | -2.102 | **0.036** | -0.101 | -0.004 |
| **Sources:Threshold** | 0.0045 | 0.004 | 1.188 | 0.235 | -0.003 | 0.012 |
| **Non-spatialized:Threshold** | 0.116 | 0.042 | 2.789 | **0.005** | 0.035 | 0.198 |
| **Sources:Non-spatialized:Threshold** | -0.0129 | 0.006 | -2.062 | **0.039** | -0.025 | -0.001 |

**Table 0.1** Estimates of each fixed effect in the full logistic regression model on performance for experiment 1 as a function of, spatialization, the number of sources and high frequency hearing threshold. $R^2$ = 0.018; Df = 7.

| | | | | | CI 95% | |
|---|---|---|---|---|---|---|
| Fixed effects | Estimate | Standard error | Z | p value | Lower | Upper |
| **Intercept** | 2.152 | 0.734 | 2.931 | **0.003** | 0.713 | 3.590 |
| **Sources** | -0.0160 | 0.112 | -0.142 | 0.887 | -0.236 | 0.204 |
| **Threshold** | 0.0641 | 0.033 | 1.913 | 0.056 | -0.002 | 0.130 |
| **Sources:Threshold** | -0.0084 | 0.005 | -1.687 | 0.092 | -0.018 | 0.001 |

**Table 0.2** Estimates of each fixed effect in the logistic regression model on performance for experiment 1 on just the non-spatialized condition as a function of the number of sources and high frequency hearing threshold. $R^2$ = 0.008; Df = 3.

| Fixed effects | Estimate | Standard error | Z | p value | CI 95% Lower | Upper |
|---|---|---|---|---|---|---|
| **Intercept** | 5.763 | 0.752 | 7.665 | **< 0.001** | 4.289 | 7.236 |
| **Sources** | -0.466 | 0.112 | -4.148 | **< 0.001** | -0.686 | -0.246 |
| **Threshold** | -0.0522 | 0.025 | -2.102 | 0.036 | -0.003 | -0.004 |
| **Sources:Threshold** | 0.0045 | 0.004 | 1.188 | 0.235 | -0.025 | 0.012 |

**Table 0.3** Estimates of each fixed effect in the logistic regression model on performance for experiment 1 on just the spatialized condition as a function of the number of sources and high frequency hearing threshold. $R^2 = 0.028$; Df = 3.

| Location 1 | Location 2 | p-value |
|---|---|---|
| Front | Right | 0.763 |
| Front | Back | **0.006** |
| Front | Left | 1.000 |
| Front | Top | **0.013** |
| Right | Back | **0.010** |
| Right | Left | 1.000 |
| Right | Top | **0.009** |
| Back | Left | **0.009** |
| Back | Top | 1.000 |
| Left | Top | **0.031** |

**Table 0.4** Table of the pairwise comparisons and associated p-values on the reaction time across participants as function of appearing source location.

| | | | | | CI 95% | |
|---|---|---|---|---|---|---|
| Fixed effects | Estimate | Standard error | Z | p value | Lower | Upper |
| ***Front*** | | | | | | |
| **Intercept** | 0.681 | 0.065 | 10.42 | < **0.001** | 0.553 | 0.809 |
| **Sources** | 0.0171 | 0.011 | 1.611 | 0.107 | -0.004 | 0.038 |
| ***Right*** | | | | | | |
| **Intercept** | 0.664 | 0.076 | 8.728 | < **0.001** | 0.515 | 0.813 |
| **Sources** | 0.0178 | 0.012 | 1.440 | 0.150 | -0.006 | 0.042 |
| ***Back*** | | | | | | |
| **Intercept** | 0.645 | 0.082 | 7.816 | < **0.001** | 0.483 | 0.806 |
| **Sources** | 0.0338 | 0.013 | 2.502 | **0.012** | 0.007 | 0.060 |
| ***Left*** | | | | | | |
| **Intercept** | 0.686 | 0.074 | 9.213 | < **0.001** | 0.540 | 0.832 |
| **Sources** | 0.0146 | 0.012 | 1.202 | 0.229 | -0.009 | 0.038 |
| ***Above*** | | | | | | |
| **Intercept** | 0.494 | 0.086 | 5.729 | < **0.001** | 0.325 | 0.663 |
| **Sources** | 0.0573 | 0.014 | 4.087 | < **0.001** | 0.030 | 0.085 |

**Table 0.5** Estimates of each fixed effect in the linear regression model on reaction time for experiment 2 for each target location (shown in bold italic) as a function of the number of sources and high frequency hearing threshold. Model fit summary: Front: $R^2$ = 0.0051; Df = 1; Right: $R^2$ = 0.0041; Df = 1; Back: $R^2$ = 0.013; Df = 1; Left: $R^2$ = 0.0029; Df = 1; Above: $R^2$ = 0.033; Df = 1.
